# PPARα genotype impacts bone quality and perfluorooctanoic acid-induced effects

**DOI:** 10.64898/2026.10.09.758097

**Authors:** Maribel O. Carpenter, Jennifer J. Schlezinger

## Abstract

Per- and polyfluoroalkyl substances (PFAS) are significant environmental contaminants that cause a number of adverse human health effects. Studies increasingly link exposure to PFAS with disruptions in osteoblast and osteoclast differentiation and function, reduced bone mineral density (BMD), and negative impacts on skeletal structure. *In vivo* studies on the effects of PFAS on bone health remain limited, with none investigating the impacts in mice expressing human peroxisome proliferator activated receptor α (PPARα). Given PFAS’s ability to interact with the peroxisome proliferator activated receptor family and their importance in bone health, we present work investigating the effects of perfluorooctanoic acid (PFOA) exposure and PPARα genotype (mouse, human or null) on bone quality across two exposure periods (6 and 14 weeks) in mice fed a human relevant diet. In both exposure scenarios, the serum PFOA concentrations were ≈ 50 μg/mL, similar to a highly exposed fluorochemical worker. We began by establishing the liver phenotype across exposures and genotypes. Across genotypes and exposure timing, PFOA increased liver to body weight ratios, but the contribution of increased triglycerides was reduced at 14 weeks. Exposure to PFOA increased hepatic PPARα-dependent gene expression (*Acox*) in both sexes and exposure periods, but only in mPPARα mice. The effect of PFOA on hepatic CAR- and PXR-dependent gene expression (*Cyp2b10* and *Cyp3a11*, respectively) was more complex; with sex-, genotype-, and exposure timing-dependent differences. Trabecular and cortical bone parameters were not impacted by PFOA treatment, but genotype was a significant factor across sexes and exposure periods. mPPARα mice had higher trabecular tissue mineral density at 9 and 20 weeks of age than hPPARα mice, but the lower trabecular bone volume evident at 9 weeks of age resolved at 20 weeks of age. mPPARα mice had higher cortical tissue mineral density and bone volume than hPPARα and PPARα null mice. Analysis of serum markers of osteoblast and osteoclast activity following 14 weeks of exposure to PFOA revealed a PPARα-dependent decrease in PINP but no effect on CTX. Our results reveal the importance of PPARα genotype for bone quality parameters and emerging evidence of the effect of PFOA on osteoblast activity. Given the unusual mouse bone quality model (sv/129 background), this work underscores the need for future in vivo research shifts to investigate the effects of PFAS on in bone quality.

## Introduction

Osteoporotic fractures are a leading cause of morbidity worldwide. Global deaths and disability-adjusted life years attributable to low bone density have doubled in the past two decades, an increase that cannot solely be explained by survival to older age (Lees et al., 1993; Sanchez-Riera et al., 2014). The annual incidence of osteoporotic fractures is greater than the incidence of heart attack, stroke and breast cancer combined in the United States (Shuler et al., 2012). Maximizing peak bone mass in adolescence is critical to reduce osteoporosis risk in old age (Hui et al., 1990; Klibanski et al., 2001). 40-60% of total adult bone mass is accrued during puberty and reductions in peak bone mass can increase the risk for osteoporosis; one model estimates an increase in peak bone mass as small as 10% predicts a 13 year delay in osteoporosis (Hereford et al., 2024; Hernandez et al., 2003). Thus, understanding the factors that impede attainment of peak bone mass is essential to improving lifelong bone health.

Millions of Americans are exposed to per- and polyfluoroalkyl substances (PFAS) daily such that PFAS are nearly universally detected in the serum of the US population (Botelho et al., 2025; CDC, 2021; De Silva et al., 2021), as well as in populations worldwide (Banjabi et al., 2020). Adverse health effects of PFAS include impairment of bone health. In analyses of general populations, serum PFAS concentrations have been associated with lower bone mineral density (Botelho et al., 2025; Cluett et al., 2019; Fan et al., 2023; Hu et al., 2019; Khalil et al., 2016; Khosrojerdi et al., 2024; L.-Y. Y. Lin et al., 2014; P.-I. D. Lin et al., 2024; Wei et al., 2024; X. Zhao et al., 2023), increased risk of osteoporosis diagnosis (Banjabi et al., 2020; Khalil et al., 2016; Khosrojerdi et al., 2024; Xu et al., 2023), and increased fracture risk (Di Nisio, De Rocco Ponce, et al., 2020). Associations between PFAS serum level and reduced bone quality have been reported for both women (Banjabi et al., 2020; Hu et al., 2019; Khalil et al., 2016; L.-Y. Y. Lin et al., 2014) and men (Di Nisio, De Rocco Ponce, et al., 2020; Hu et al., 2019; Khalil et al., 2016). Associations between serum PFAS concentrations and reduced bone quality also have been observed in children (Buckley et al., 2021; Carwile et al., 2022; Cluett et al., 2019; Højsager et al., 2023; Jeddy et al., 2018; Khalil et al., 2018; Rokoff et al., 2024; Sigvaldsen et al., 2025; Xiong et al., 2022). Further, reduced bone formation is a critical health effect underlying regulatory PFAS health advisories and drinking water guideline in the U.S. (USEPA, 2016).

The weight of evidence is building to support the hypothesis that PFAS can impair bone health. **1)** PFAS are present in human bone (Koskela et al., 2017; Perez et al., 2013). **2)** While PFAS are charged at physiological pH and must be actively transported into cells at concentrations found in humans (Kimura et al., 2017). osteoblasts express members of the OATP family of the transporter proteins known to transport PFAS into cells.(Di Nisio, De Rocco Ponce, et al., 2020; Kudo et al., 2002; Monjo et al., 2010; Weaver et al., 2010; W. Zhao et al., 2017). **3)** In mouse models (**Table S1**), *in utero* exposure to perfluorooctanoic acid (PFOA) caused reduced ossification in term fetuses. (Lau et al., 2006) I*n utero*/lactational exposure to 6:2 Cl-PFESA resulted in shortened femur length and reduce bone quality parameters at PND21(Feng et al., 2024) and to PFOA resulted lower mineral density in the tibia and a larger medullary area in the femur and tibia at 17 months of age.(Koskela et al., 2016) **4)** In exposures spanning puberty, PFAS reduced bone length and measures of trabecular bone quality.(Wang et al., 2024a) **5)** In chronic adult exposures, perfluoro (2-methyl-3-oxhexanoic) acid (HFPO-DA) and perfluorohexane sulfonic acid (PFHxS) caused reductions in trabecular bone volume, which were thought to be the major contributors to the effect of a mixture also including PFOA, perfluorononanoic acid (PFNA), and perfluorooctane sulfonic acid (PFOS).(Garmo et al., 2025a) **6)** PFAS have been shown to increase human and mouse primary osteoclast resorptive activity and number and size of osteoclasts.(Garmo et al., 2025a; Koskela et al., 2016, 2017; Liu et al., 2019) However, *in vitro* studies of PFAS-induced effects in osteoblasts have produced mixed results.(Di Nisio, Rocca, et al., 2020; Koskela et al., 2016, 2017; Liu et al., 2019; Pan et al., 2019a)

Peroxisome proliferator activated receptor α (PPARα) has recently been identified as a contributor to bone homeostasis and quality (Chougule et al., 2023). PPARα also is a well-documented contributor to PFAS-driven effects in mice expressing either mouse or human PPARα (Nielsen et al., 2026; Rosen et al., 2017). Therefore, we tested the hypothesis that PFAS can compromise bone quality via interaction with PPARα. PPARα is well-known for its species-specificity (Takacs & Abbott, 2007) (Corton et al., 2018). The present study investigates the effect of PFOA exposure on trabecular and cortical bone in mice expressing human PPARα, in comparison to mice expressing mouse PPARα and PPARα null mice.

## Materials and Methods

### Materials

Perfluorooctanoic acid (95% purity, cat. #171468) was purchased from Sigma-Aldrich (St. Louis, MO). All other reagents were sourced from Thermo Fisher Scientific (Waltham, MA), unless otherwise noted.

### In vivo exposures

All animal studies were approved by the Institutional Animal Care and Use Committee at Boston University and performed in an American Association for the Accreditation of Laboratory Animal Care accredited facility (Animal Welfare Assurance Number: A3316-01). Male and female, humanized PPARα mice (hPPARα) were generated from mouse PPARα null, human PPARα heterozygous breeding pairs (generously provided by Dr. Frank Gonzalez, NCI)(Q. Yang et al., 2008). Genotyping for mouse and human PPARα was carried out by Transnetyx (Cordova, TN). The expression level of hPPARα in liver was confirmed by RT-qPCR. Wildtype 129S4/SvJae mice (009104, 6-week exposure) and 129S1/SvlmJ mice (002448, 14-week exposure) were purchased from Jackson Laboratory (Bar Harbor, ME). The 6-week exposure did not originally include a wildtype mouse, as such the strains differ because we were unsure of the best match to the original sv/129 strain into which the hPPARα transgene was introduced (Lee et al., 1995). We ultimately found a study suggesting that the Jae strain was a better match, as such the Jae strain was utilized for the 6 week exposure where a WT was not originally included (Akiyama et al., 2001).

### *6-week* PFOA exposure

Exposure of the hPPARα and PPARα null mice to PFOA has been previously described (Schlezinger, Puckett, Nielsen, et al., 2020). Here, we generated new data only in wildtype (mPPARα) mice for the 6-week exposure. At weaning or on arrival, mice were provided a custom diet based on the “What we eat in America (NHANES 2013/2014)” analysis for what 2-19 year old children and adolescents eat (Research Diets, New Brunswick, NJ)(USDA, 2018). The composition as % energy intake of the “What we eat in America” adolescent diet is 51.8% carbohydrate, 33.5% fat, and 14.7% protein (**Table S2**). Fats are in the form of soybean oil, lard and butter, with cholesterol at 0.07 mg/kcal. Vehicle (Vh) and treatment water were prepared from NERL High Purity water (23-249-589, Thermo Fisher Scientific), which is prepared using reverse osmosis and carbon filtering). A concentrated stock solution of PFOA (1×10^-2^ M) was prepared in NERL water and then diluted in NERL water containing 0.5% sucrose. Mice were administered vehicle (0.5% sucrose) drinking water or PFOA (8 μM, 3.3 mg/L) drinking water *ad libitum* for 6-7 weeks.

### *14-week* PFOA exposure

Exposure of the mPPARα, hPPARα and PPARα null mice has been previously described (Nielsen et al., 2025). At 6-7 weeks of age, mice were provided a custom diet based on the “What we eat in America (NHANES 2015/2016)” analysis for 20+ year old adults (Research Diets, New Brunswick, NJ)(USDA, 2021). The composition as % energy intake of the “What we eat in America” adult diet is 47.3% carbohydrates, 36.7% fat, and 16.0% protein (**Table S2**). Fats are in the form of soybean oil, lard and butter, with cholesterol at 0.25 mg/kcal. Drinking water was prepared as described above. Mice were administered vehicle (0.5% sucrose) drinking water or PFOA (3.4 μM, 1.4 mg/L) drinking water *ad libitum* for 14 weeks. There was select serum available from mice exposed to PFOA at 6.2mg/L in drinking water– this serum was only utilized in the PINP and CTX serum quantification and is classified as “PFOA High” vs the typical PFOA exposure of 1.4mg/L which is designated as “PFOA Low”.

### Experimental husbandry

Food and water consumption were determined on a per cage basis each week. Body weight was measured weekly. Mice were analyzed for body composition (total body fat mass, lean mass, water) using an EchoMRI700 (EchoMRI LLC, Houston, TX), fasted for 6 hours and then euthanized. Blood was collected via cardiac puncture and serum separated. Livers were collected from each mouse and weighed. Aliquots of liver for gene expression and triglyceride analyses were flash frozen in liquid nitrogen and stored at -80°C. The left femur was dissected, wrapped in phosphate buffered-saline soaked gauze, placed in a screw-top 5 ml tube and stored at -80°C. The left tibia was dissected, fixed in 10% neutral buffered formalin for 48 hours and then stored in 70% ethanol at 4°C. The right femur and tibia were dissected, with attention paid to removing all muscle tissue, flash frozen in liquid nitrogen and stored at -80°C.

### PFAS analysis

Serum PFOA concentration was determined in [N = 3-5] samples pooled by cage per sex and exposure group. Serum PFOA concentrations were measured by LC-MS/MS using method MLA-042 Rev 09 (SGS AXYS Analytical Services Ltd., Sydney, B.C., Canada).

### Liver triglyceride analysis

Liver triglyceride content was measured using methods that have been described previously (Norris et al., 2003). We prepared a saponified neutralized liver extract and assessed triolein content using free glycerol reagent (Sigma F6428). A standard curve was generated with glycerol standard solution (Sigma G7793). Absorbance was determined at 540 nM on a Synergy2 plate reader (Biotek Inc, Winooski VT, USA). Triolein was converted to triglyceride and divided by wet liver weight to derive liver lipid concentration in mg triglycerides/g liver.

### Micro-Computed Tomography (micro-CT)

Sample analyses were performed according to guidelines outlined by M.L. Bouxsein (Bouxsein et al., 2010). Samples were scanned using a Scanco micro-CT 40 system (Scanco Medical; Brütisellen, Switzerland) using power, current, and integration time of 70 kVP, 113 mA, and 200 ms, respectively. Gaussian filtering (sigma = 0.8, support = 1) was used for reducing background noise. To analyze trabecular bone in the distal femur, the distal metaphysis was scanned at a nominal resolution of 8 µm/voxel, and the region of interest analyzed began at 0.08 mm proximal to the growth plate and extended 0.8 mm proximally. The trabecular compartment was automatically segmented from the cortical shell (Scanco Medical). Cortical bone was analyzed from 12 µm/voxel scans of a 0.6 mm-tall region at the mid-diaphysis. For trabecular bone, sample-specific thresholding was used (mean value: 483.1 mg HA/cm^3^) with thresholds automatically determined by an iterative method (Scanco Medical). Cortical bone was also evaluated at a sample-specific threshold (mean value: 813.9 mg HA/cm^3^). Tissue mineral density was calculated with the aid of a standard curve obtained from a scan of a hydroxyapatite phantom consisting of five different mineral densities (Scanco Medical).

### Histology

Bones were prepared for histological analysis, imaged and analyzed by the Bone Histomorphometry Core Laboratory at the MD Anderson Cancer Center (Houston, Tx). Tibiae were decalcified in 0.5 M sodium EDTA solution (Sigma-Aldrich, E5134) for a week with gentle stirring at 4°C, the EDTA solution was replaced, and the tissue was decalcified for an additional week. The tibiae were transferred and processed for dehydration and paraffin infiltration with the Leica HistoCore PELORIS 3 using standard processing protocol. Then tibiae were embedded in paraffin, and 4µm slices were sectioned using the Leica HistoCore Arcadia Embedding station and NANOCUT rotary microtome. Slices were stained with hematoxylin and eosin (H&E), tartrate-resistant acid phosphatase (TRAP). Sections were digitized using a Nikon Eclipse Ti2 inverted microscope. The images were acquired in BIF format using the “Once/Field” focus pattern setting and a 20x objective magnification was utilized.

### Gene expression analyses

Livers were homogenized in TRI Reagent. Right tibia and femur were crushed by mortar and pestle in TRI Reagent. Total RNA was extracted and genomic DNA was removed using the Direct-zol RNA Microprep (bone) or Miniprep (liver) Kit (Zymo Research, Orange, CA). cDNA was synthesized from total RNA using the iScript™ Reverse Transcription System (BioRad, Hercules, CA). All qPCR reactions were performed using the PowerUp™ SYBR Green Master Mix (Thermo Fisher Scientific, Waltham, MA). The qPCR reactions were performed using a StepOnePlus Real-Time PCR System (Applied Biosystems, Carlsbad, CA). The primer sequences and annealing temperatures are provided in **Table S3**. Relative gene expression was determined using the Pfaffl method to account for differential primer efficiencies (Pfaffl, 2001), using the geometric mean of the Cq values for beta-2-microglobulin (*B2m*), GAPDH (*Gapdh*), and 18sRNA (*R18s*) for liver and *B2m* and *R18s* for bone. The average Cq value from independent samples was used as the reference point. Data are reported as “Relative Expression.”

### Serum Protein Analysis

CTX-I and PINP serum concentrations were determined using Immunodiagnostic Systems (IDS) Rat/Mouse PINP EIA (REF:AC-33F1) and IDS RatLaps (CTX-I) EIA (REF:AC-06F1). All plates were prepared with calibration values for the standard curve, control samples with known concentration, and analytical samples. All analytes were run in duplicates when possible and samples above appropriate coefficient of variations (CV%’s) as determined by the lab performance data were excluded. Given limited sample availability for each genotype and treatment group sexes were combined for these analyses.

### Statistical analyses

Data are presented as data points from individual mice or as means ± standard deviation (SD). Mice were considered hPPARα positive if they were either homozygous or heterozygous. All statistical tests were run in Prism 11 (GraphPad, Boston, MA). Within sex and genotype, statistical significance was determined by unpaired, two-tailed t-test. ROUT analyses were utilized to identify outliers in qPCR and serum quantification of PINP and CTX (Q = 0.1%).

Statistical significance was evaluated at an α= 0.05 for all analyses.

## Results

### Serum PFOA concentrations

Serum PFOA concentrations after the 6- and 14-week exposure are shown in **Figure 1**. Exposure to 3.3 mg PFOA/L in drinking water for 6 weeks and to 1.4 mg PFOA/L in drinking water for 14 weeks resulted in similar PFOA serum concentrations in most groups. A significant sex-difference was detected in the 14-week exposure, with males having lower serum PFOA concentrations than females (Two-Factor ANOVA, sex p = 0.0189, treatment p < 0.0001, sex x treatment p = 0.0189). This difference is consistent with the observation that females drank more water than males (Nielsen et al., 2025). Given these differences, all results are stratified by sex to maintain consistency between the two exposure groups.

**Figure 1.**
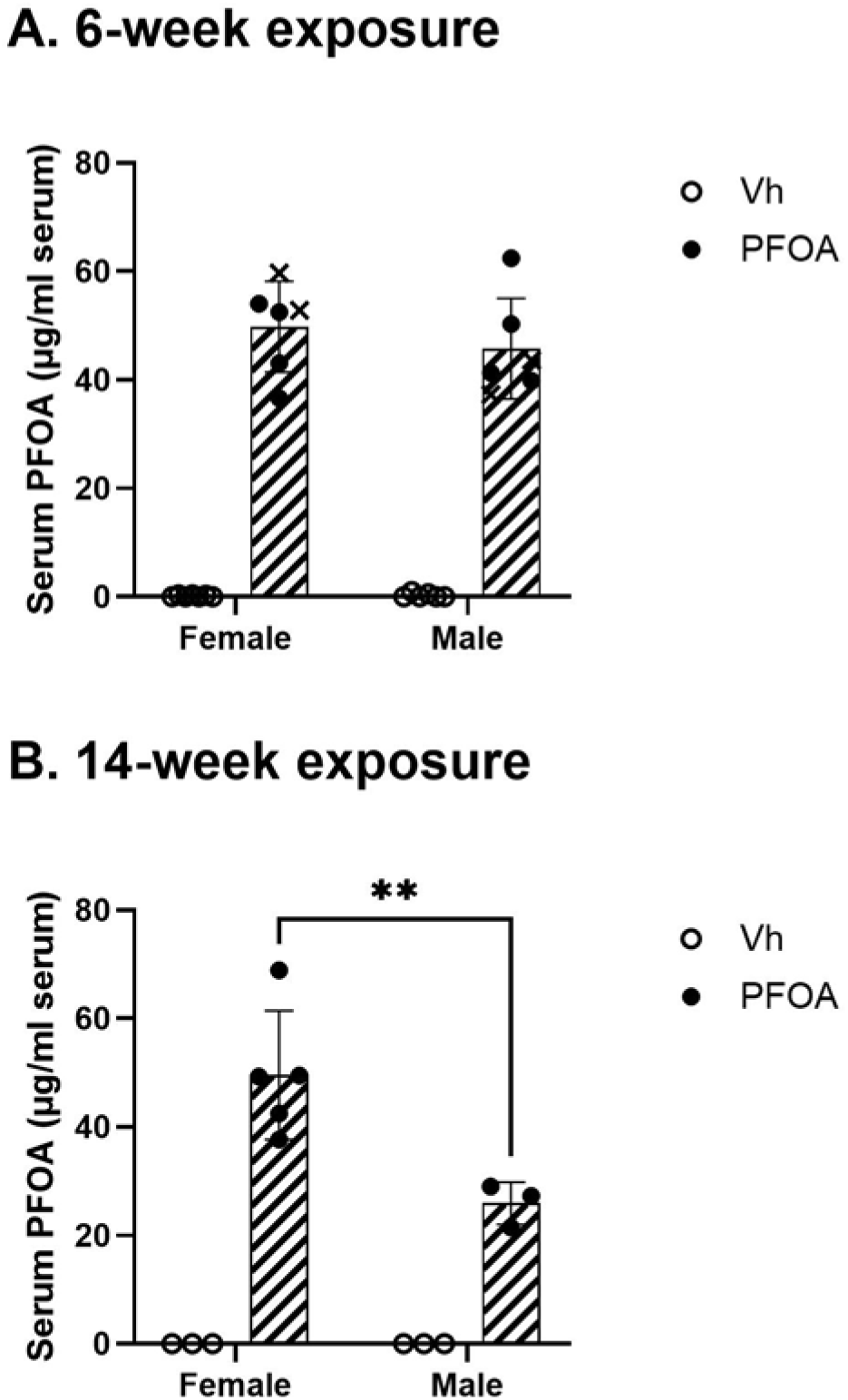
Serum PFOA concentrations in vehicle and exposed mice. **(A)** Three to four-week-old female and male hPPARα, PPARα null and WT mice were treated with either vehicle (Vh, NERL water with 0.5% sucrose) or PFOA (8 µM (3.3 mg/L)) in NERL water with 0.5% sucrose) as drinking water for 6 weeks. During treatment, the mice were fed an adolescent American Diet. Data from hPPARα and PPARα null mice were previously published (Schlezinger, Puckett, Nielsen, et al., 2020). Data from the mPPARα cohorts are indicated with an X as they were collected after the initial experiment. N = 3-6. **(B)** Six to seven-week-old female and male hPPARα, PPARα null and WT mice were treated with vehicle (Vh, NERL water with 0.5% sucrose) or PFOA (3.4 μM (1.4 mg/L) in NERL water with 0.5% sucrose) for 14 weeks. During treatment, the mice were fed an adult American Diet. Data were previously published (Nielsen et al., 2025). PFAS concentrations were determined by LC-MS/MS. Data are from individual mice or pooled from mice in the same cage, with the mean indicated the bar and the SD indicated by the error bars. N = 3-6. Effects of sex and exposure were tested using a two-way ANOVA. ** p<0.01 (Šídák’s test).

### Effect of PFOA on liver physiology

We further verified that a toxicologically effective dose of PFOA was achieved by analyzing liver tissue. Importantly, studies rarely have investigated changes in liver morphology in response to differing times of PFAS exposure. Therefore, we began by analyzing well-known effects of PFOA in liver over both exposure periods to inform our analysis of effects in bone.

After 6 weeks of exposure, PFOA significantly increased liver to body weight ratio in both sexes and all genotypes (PPARα null, hPPARα and mPPARα)(**Figure 2A,C**). Similarly, at 14 weeks of exposure, PFOA significantly increased liver to body weight ratios in female mice of all genotypes (**Figure 2A,C**). However, in males, PFOA significantly increased liver to body weight ratios in mice expressing hPPARα or mPPARα but not in PPARα null mice (**Figure 2**). The effects of PFOA on liver triglyceride content across genotypes and time were more complex (**Figure 2B,C**). After 6 weeks of exposure, PFOA increased liver triglycerides in hPPARα and mPPARα female mice but increased liver triglycerides only in hPPARα male mice. PFOA did not have a significant effect on liver triglycerides in PPARα null mice. After 14 weeks of exposure, PFOA did not significantly increase liver triglycerides in either sex or genotype except for a PFOA-induced decrease in liver triglycerides in PPARα null female mice. However, after 14 weeks all the mice showed a marked increase in liver triglycerides compared to the 6-week exposure, indicating potential variation due to timing of exposure.

**Figure 2.**
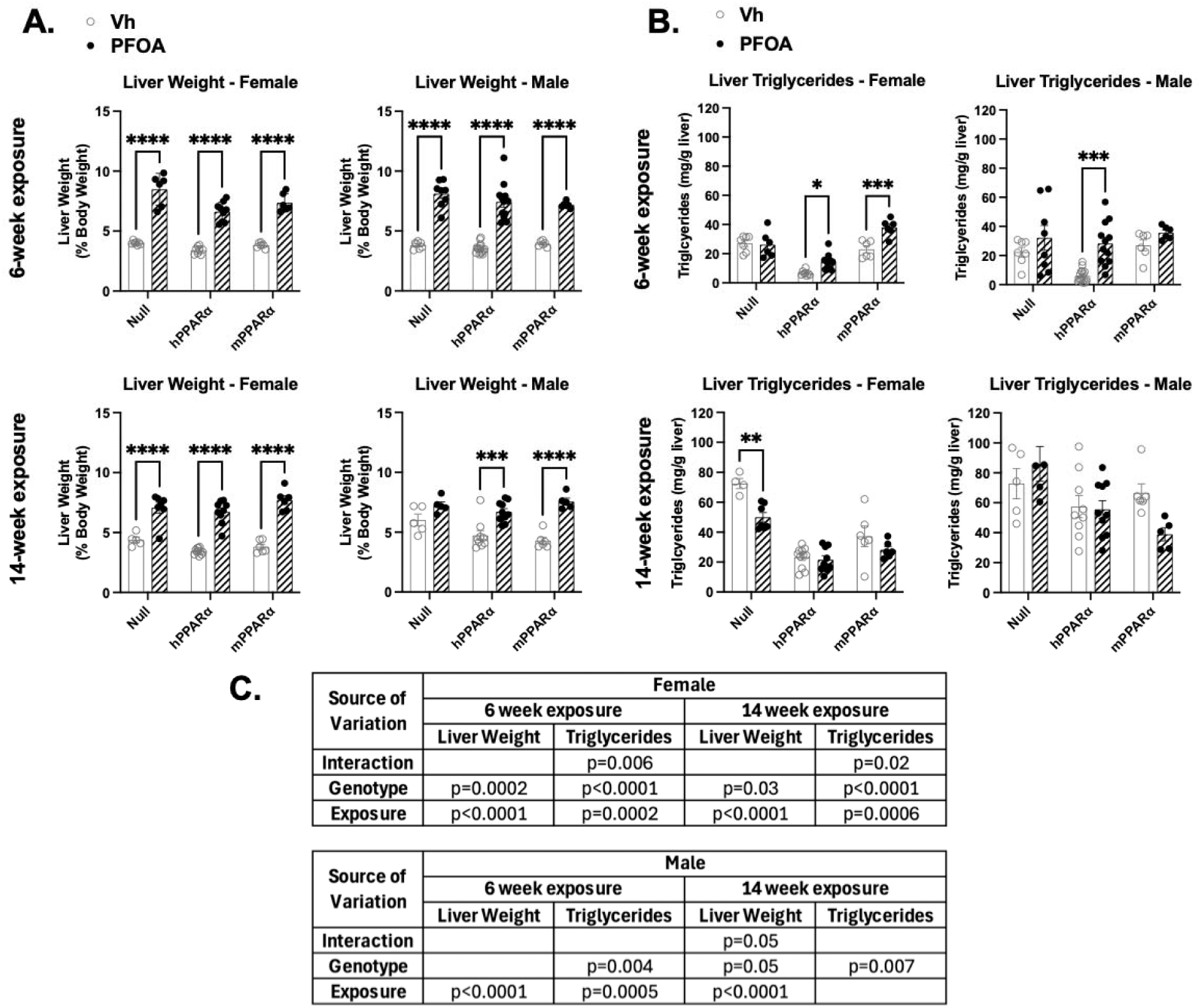
Liver phenotype in vehicle and PFOA exposed mice. Samples were analyzed from two experimental scenarios, as described in Figure 1. **(A)** Liver to body weight ratios were determined from weights taken at euthanasia. **(B)** Total triglyceride concentrations were determined from measuring triolein content in saponified liver extracts. **(C)** Results of two-way ANOVA. Data points are from individual mice, with the mean indicated the bar and the SD indicated by the error bars. N = 6-9. ** p<0.01, *** p<0.001, **** p<0.0001 (Tw-way ANOVA, Šídák’s test).

### Effects of PFOA on liver gene expression

PPARα, CAR and PXR pathways are well known to be upregulated by PFOA (Abe et al., 2017; Behr et al., 2020; Bjork et al., 2011; Louisse et al., 2020; Murase et al., 2023). Target gene expression for each nuclear receptor was analyzed to assess PFOA-induced activation. *Acox*, a PPARα target gene was upregulated by PFOA in mPPARα and hPPARα mice, regardless of sex or timing of exposure but this difference was only significant for mPPARα (**Figure 3**). After 6 weeks of exposure, *Cy2b10*, a CAR target gene, was more strongly upregulated by PFOA in hPPARα and PPARα null mice than in mPPARα mice, in both sexes (**Figure 3**). After 14 weeks of exposure, *Cyp2b10* was only upregulated by PFOA in hPPARα for males and females and male PPARα null mice. Notably, the level of expression of *Cyp2b10* was ≈ 2x higher at 14 weeks of exposure than 6 weeks of exposure. After 6 weeks of exposure, *Cyp3a11*, a PXR target gene, was upregulated by PFOA in all but male mPPARα mice (**Figure 3**). After 14 weeks of exposure, *Cyp3a11* was upregulated by PFOA in all but female mPPARα mice. The level of expression of *Cyp3a11* was ≈ 10x lower at 14 weeks of exposure than 6 weeks of exposure.

**Figure 3.**
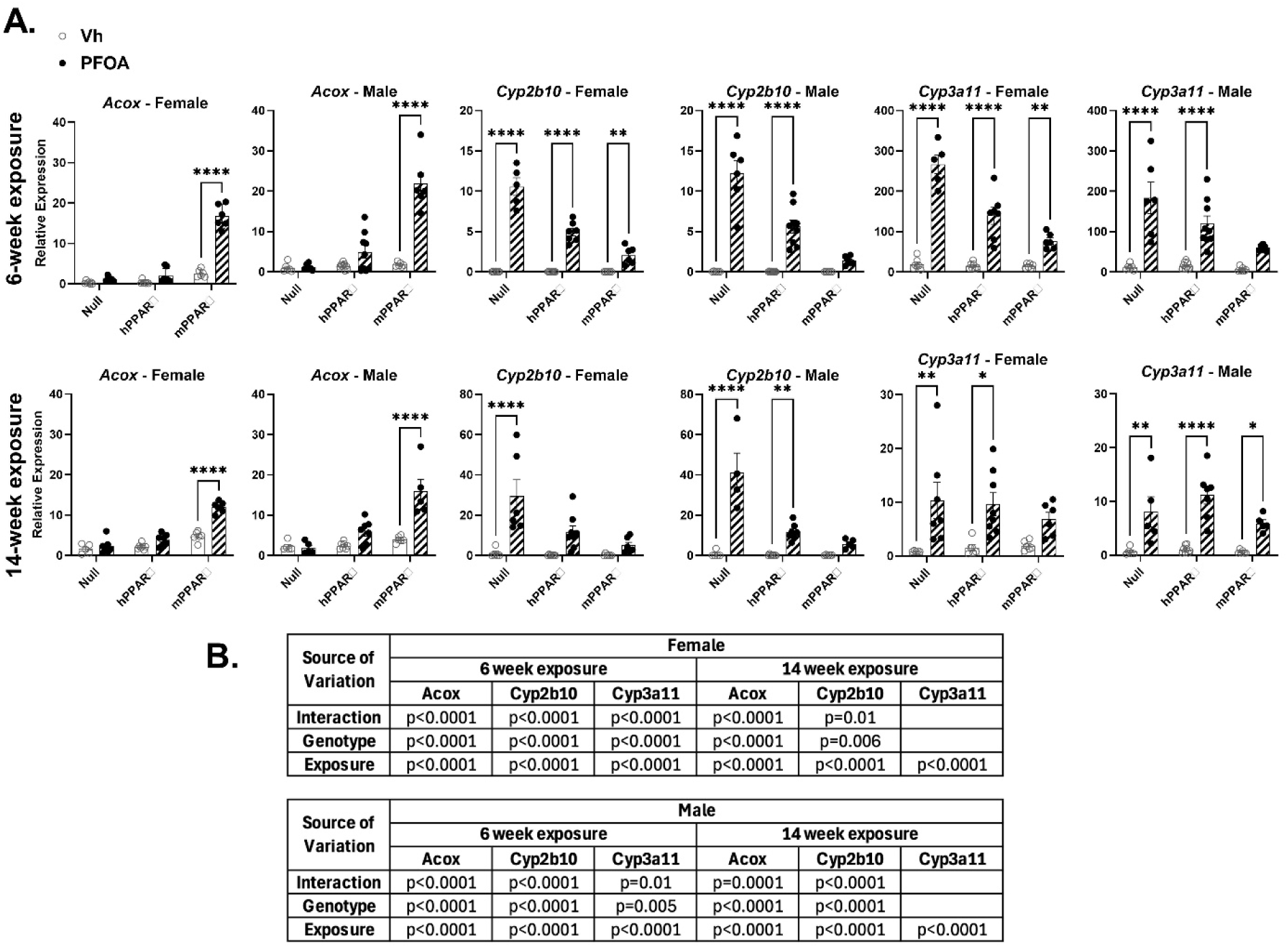
PPARα, CAR and PXR target gene expression in liver of vehicle and PFOA exposed mice. Samples were analyzed from two experimental scenarios, as described in Figure 1. **(A)** . Following isolation of RNA from liver, gene expression was determined by RT-qPCR. **(B)** Results of two-way ANOVA. Data points are from individual mice, with the mean indicated the bar and the SD indicated by the error bars. N = 6-9. *p<0.05, ** p<0.01, ****p<0.0001 (Two-way ANOVA, Šídák’s test).

### Effect of PPARα genotype on trabecular structure

Trabecular structure was analyzed using micro-CT at the distal metaphysis of the femur. No effect of PFOA on trabecular structure was observed at either timepoint (**Table S4**); therefore, data are presented and analyzed solely based on sex and genotype.

After 6 weeks on the WWEIA diet (9 weeks of age), micro-CT images show marked differences in trabecular structure in female mice between genotypes (**Figure 4A**). Data from male mice are presented in **Figure S1**. Trabecular tissue mineral density (TMD) was lower in both hPPARα and PPARα null mice than in mPPARα and the ratio of trabecular bone volume to total volume (BV/TV), connective density (Conn.D), and number (Tb.N) was significantly higher in hPPAR⍰ mice compared to mPPAR⍰ (**Figure 4B**). Furthermore, for BV/TV, Conn.D, and Tb.N PPARα null mice were numerically lower than hPPARα mice and numerically higher than mPPARα mice, although non-significant. Consistent with this, trabecular spacing (Tb.Sp) was significantly lower in hPPAR⍰ mice compared mPPAR⍰ and PPAR⍰ null mice were numerically lower than mPPAR⍰ and higher than hPPAR⍰ (**Figure 4B**). No differences were observed in trabecular thickness (Tb.Th) across genotypes. Similar patterns were observed for male mice (**Figure S1)**.

**Figure 4.**
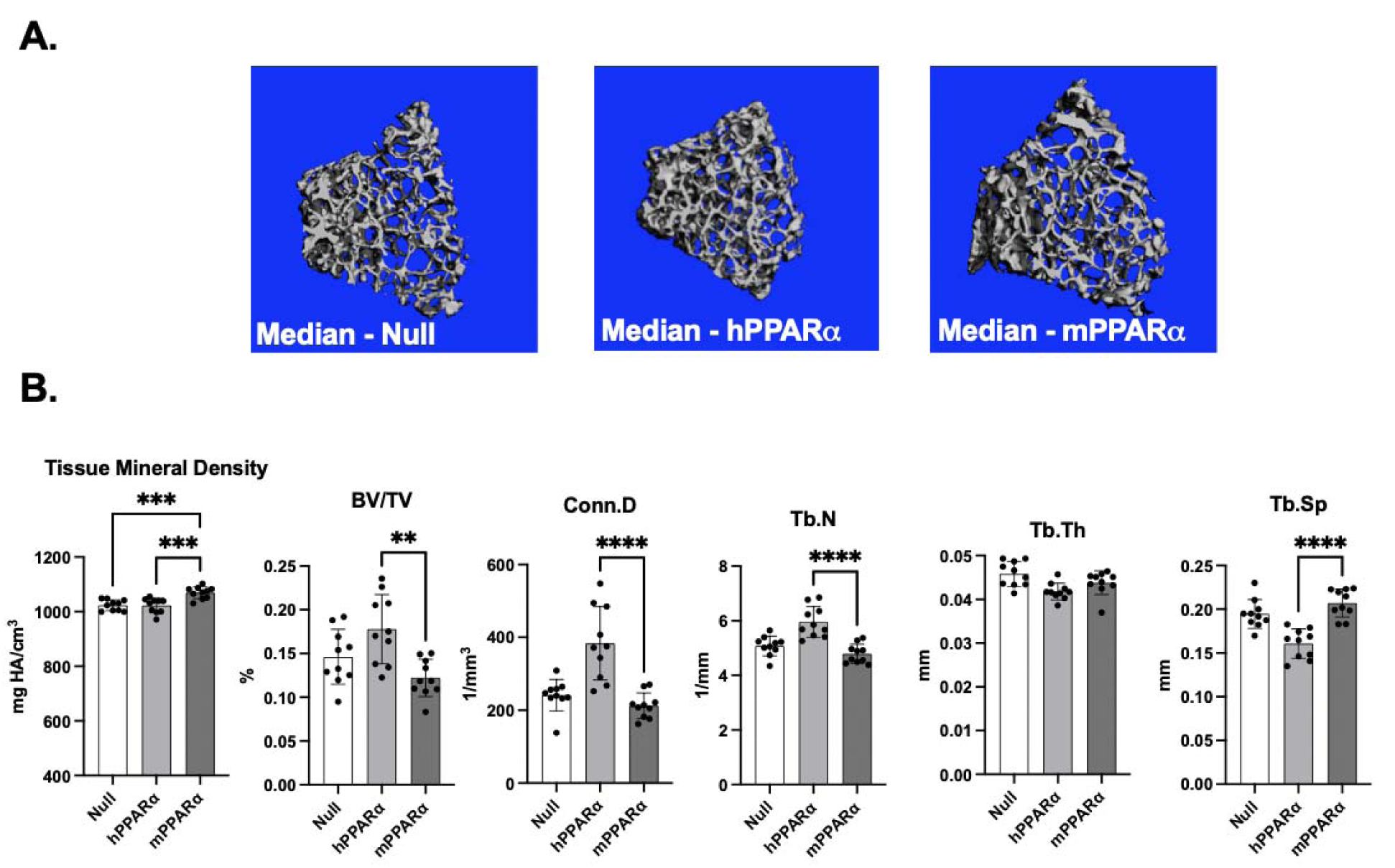
Micro-CT analysis of femur trabecular bone structure for female mice after 6-week PFOA exposure. Samples were analyzed from the 6-week exposure described in Figure 1; mice were 9-weeks of age. **(A)** Representative (i.e., bones with the median connective density for each genotype) micro-CT images of distal metaphysis of female mice. **(B)** Trabecular bone parameters of female mice. BV/TV: bone volume to total volume ratio; Conn.D: connective density; Tb.N: number of trabeculae; Tb.Th: trabecular thickness; Tb.Sp: trabecular spacing. Data points are from individual mice, with the mean indicated the bar and the SD indicated by the error bars. N = 6-9. *p<0.05, ** p<0.01, *** p<0.001, ****p<0.0001 (ANOVA, Dunnett’s test versus mPPARα).

After 14 weeks on the WWEIA diet (20 weeks of age), micro-CT images show substantial differences in trabecular structure in female mice between genotypes (**Figure 5A**). Data from mice are presented in **Figure S2.** Trabecular TMB was lower in hPPARα females than in mPPARα mice (**Figure 5B**). Trabecular BV/TV, connective density, and number were lower in PPAR⍰ null mice compared to mPPAR⍰, but hPPAR⍰ mice were more like mPPAR⍰ (**Figure 5B**). These results were only significant in male mice (**Figure S2**). Similarly, trabecular spacing was significantly higher in PPAR⍰ null mice compared to mPPAR⍰, with hPPAR⍰ mice more comparable to mPPAR⍰ mice. However, trabecular thickness was significantly lower in hPPAR⍰ and higher in PPAR⍰ null mice compared to mPPAR⍰ (**Figure 5B**). Similar patterns for trabecular spacing and thickness were observed in males (**Figure S2**).

**Figure 5.**
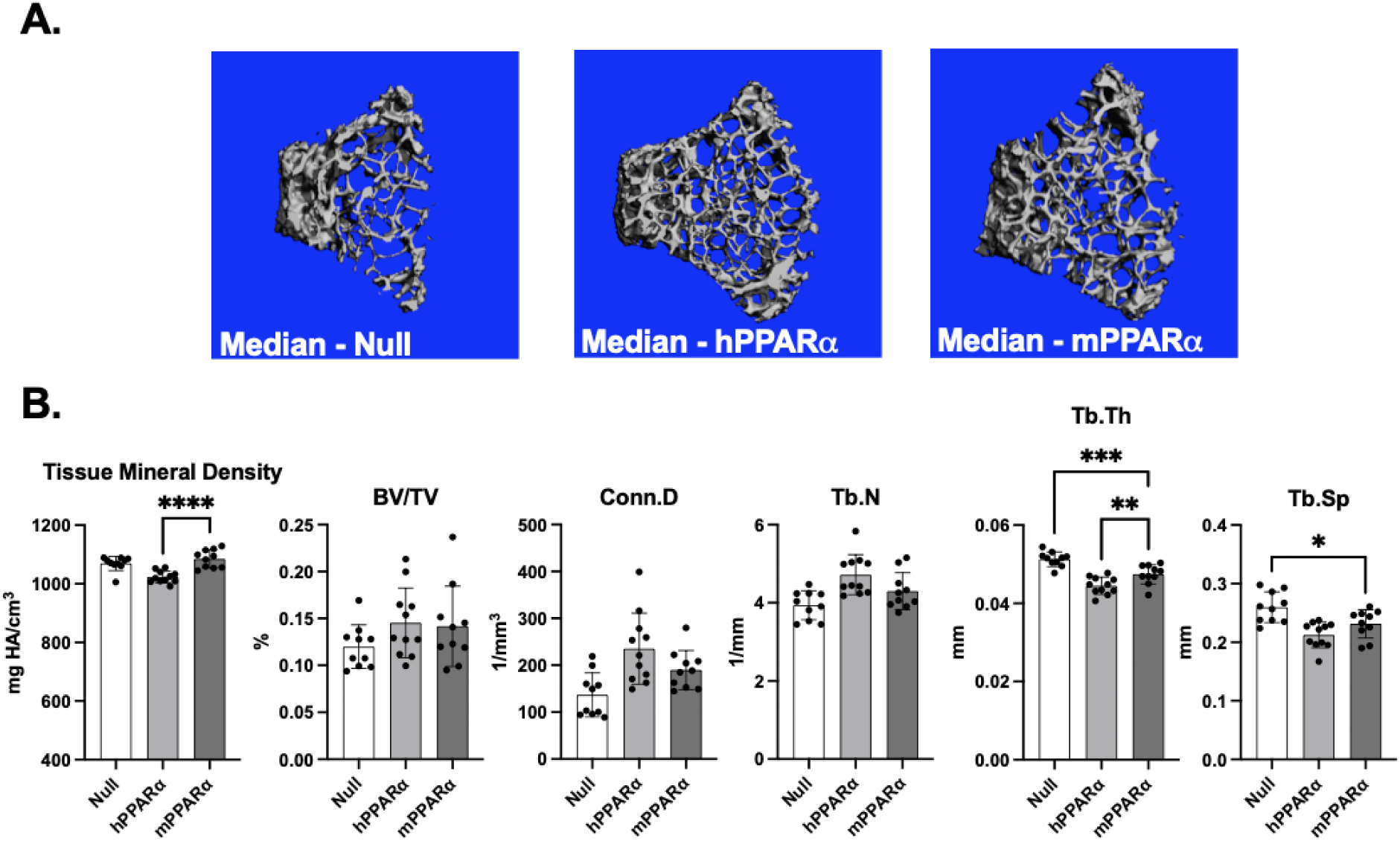
Micro-CT analysis of femur trabecular bone structure for female mice after 14-week PFOA exposure. Samples were analyzed from the 14-week exposure described in Figure 1; mice were 20-weeks of age. **(A)** Representative (i.e., bones with the median connective density for each genotype) micro-CT images of distal metaphysis of female mice. **(B)** Trabecular bone parameters of female mice. BV/TV: bone volume to total volume ratio; Conn.D: connective density; Tb.N: number of trabeculae; Tb.Th: trabecular thickness; Tb.Sp: trabecular spacing. Data points are from individual mice, with the mean indicated the bar and the SD indicated by the error bars. N = 6-9. *p<0.05, ** p<0.01, *** p<0.001, ****p<0.0001 (ANOVA, Dunnett’s test versus mPPARα).

### Effect of PPAR_α_ genotype on cortical structure

Cortical structure was analyzed using micro-CT at the mid-diaphysis of the femur. No effect of PFOA on cortical structure was observed at either timepoint (**Table S5**), therefore data are presented and analyzed solely based on sex and genotype.

After 6 weeks on the WWEIA diet (9 weeks of age), micro-CT images show moderate differences in cortical thickness in female mice between genotypes (**Figure 6A**). Data from male mice are presented in **Figure S3**. Cortical TMD, BV/TV, and thickness (Cort.Th) were significantly lower in both hPPAR⍰ and PPAR⍰ null mice compared to mPPAR⍰ (**Figure 6B**). Cortical area (Cort.Ar) was smaller in hPPAR⍰ mice compared to mPPAR⍰, and cortical marrow area (Cort.Mar.Ar) was smaller for PPAR⍰ null mice compared with mPPAR⍰. No differences were observed in total cortical area (Cort.Total.Ar). Similar trends were observed in male mice (**Figure S3**).

**Figure 6.**
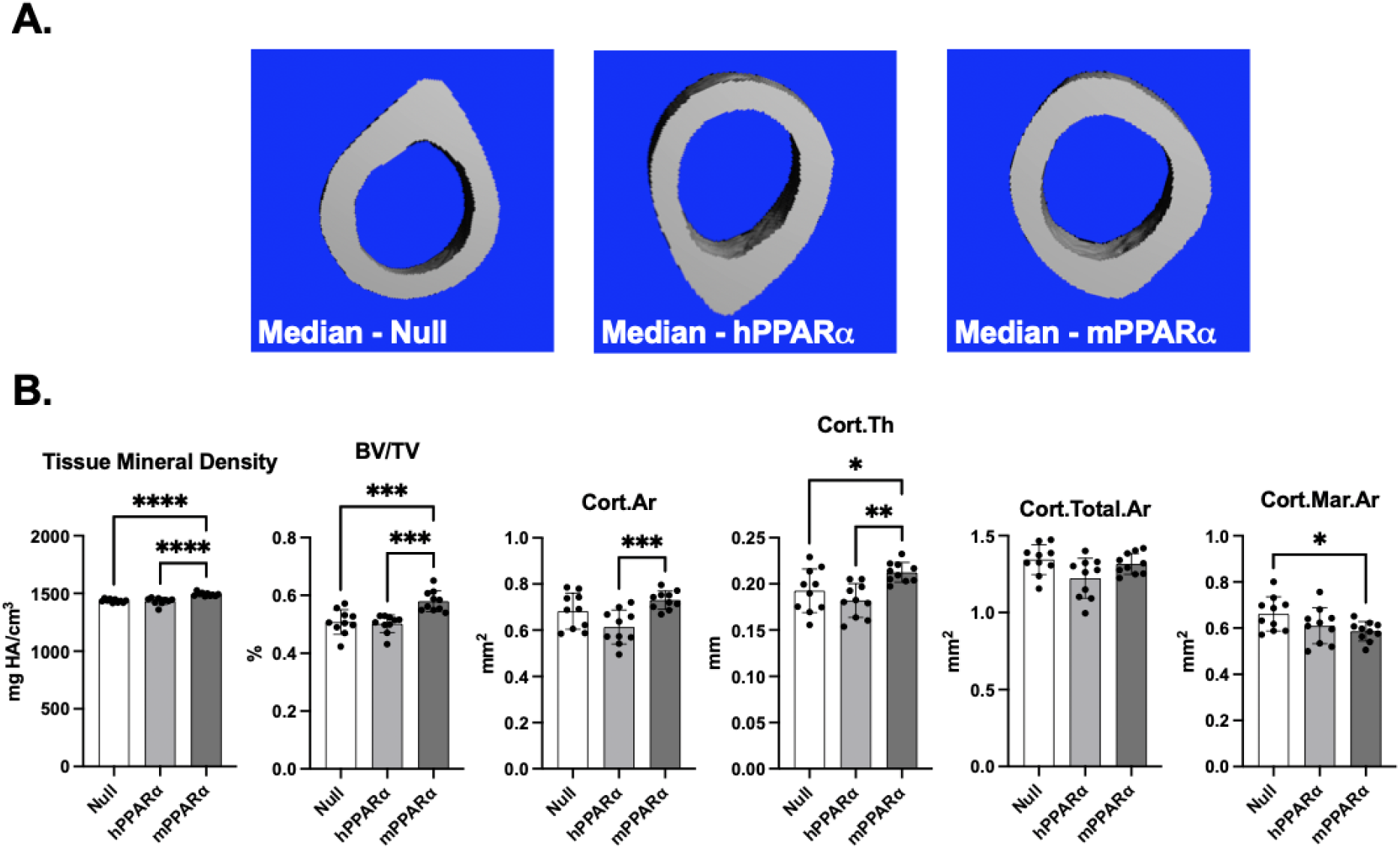
Micro-CT analysis of femur cortical bone structure for female mice after 6-week PFOA exposure. Samples were analyzed from the 6-week exposure described in Figure 1; mice were 9-weeks of age. **(A)** Representative (i.e. bones with median cortical thickness for each genotype) micro-CT images of mid-diaphysis of female mice. **(B)** Cortical bone parameters of female mice. BV/TV: bone volume to total volume ratio, Cort.Ar: total cortical area, Cor.Th: cortical thickness, Cort.Total.Area: total bone area, Cort.Mar.Ar: Cortical marrow area. Data points are from individual mice, with the mean indicated the bar and the SD indicated by the error bars. N = 6-9. *p<0.05, ** p<0.01, *** p<0.001, ****p<0.0001 (ANOVA, Dunnett’s test versus mPPARα).

After 14 weeks on the WWEIA diet (20 weeks of age), micro-CT images show consistent reductions in cortical thickness between lowest and median for each genotype (**Figure 7A**). Data from male mice are presented in **Figure S4**. Cortical TMD, BV/TV, and Cort.Th were lower for hPPAR⍰ and PPAR⍰ null mice compared to mPPAR⍰ (**Figure 7B**). Cort.Ar was lower for hPPAR⍰ only compared to mPPAR⍰. Similar trends were observed for male mice (**Figure S4**). No differences were observed for Cort.Total.Ar or Cort.Mar.Ar, except for male mice where PPAR⍰ null mice had increased Cort.Mar.Ar compared to mPPAR⍰ (**Figure 7B** and **Figure S4**). *Effect of PFOA on Osteoblasts and Osteoclasts*

**Figure 7.**
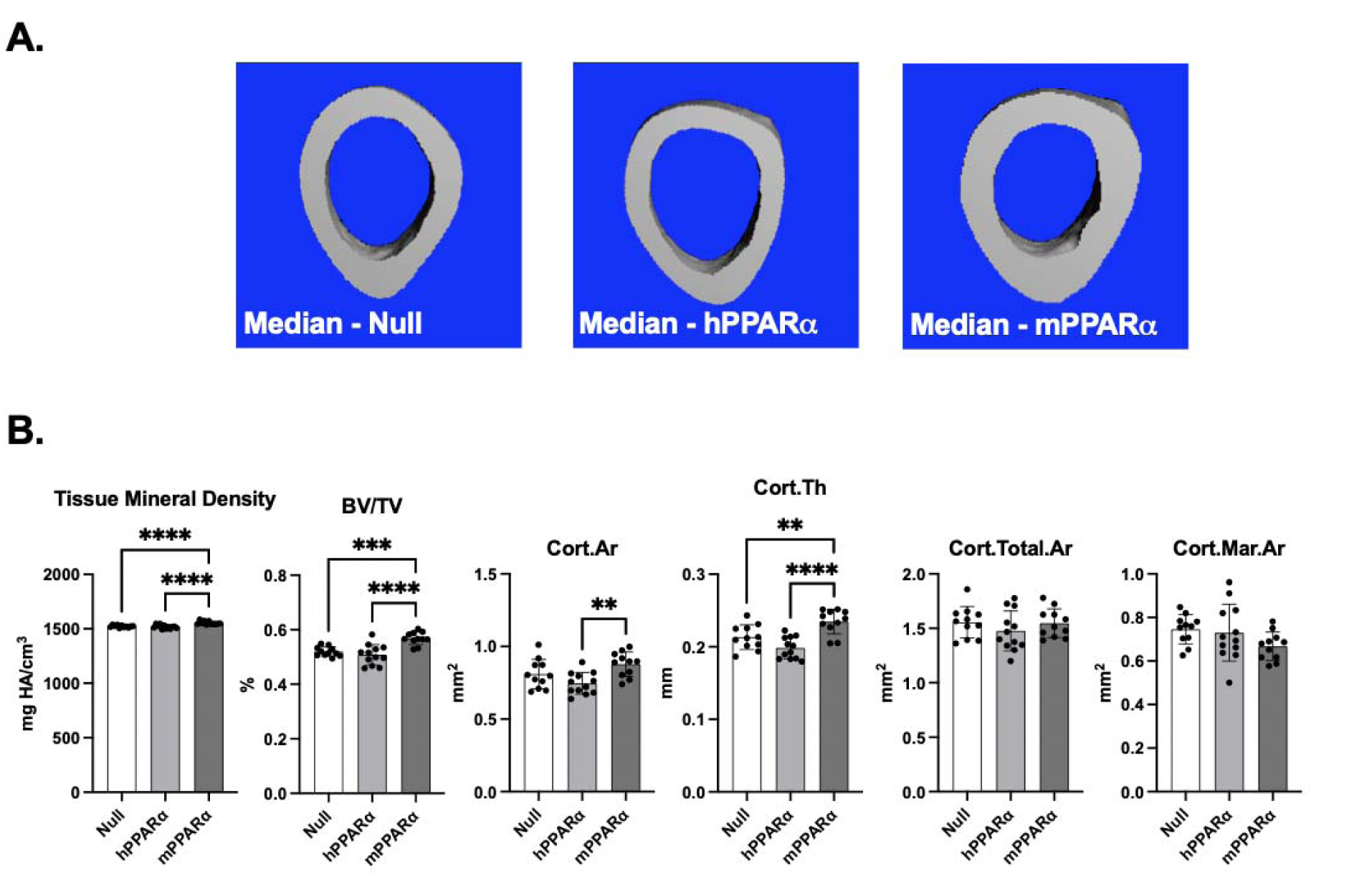
Micro-CT analysis of femur cortical bone structure for female mice after 14-week PFOA exposure. Samples were analyzed from the 14-week exposure described in Figure 1; mice were 20-weeks of age. **(A)** Representative (i.e. bones with median cortical thickness for each genotype) micro-CT images of mid-diaphysis of female mice. **(B)** Cortical bone parameters of female mice. BV/TV: bone volume to total volume ratio, Cort.Ar: total cortical area, Cor.Th: cortical thickness, Cort.Total.Area: total bone area, Cort.Mar.Ar: Cortical marrow area. Data points are from individual mice, with the mean indicated the bar and the SD indicated by the error bars. N = 6-9. *p<0.05, ** p<0.01, *** p<0.001, ****p<0.0001 (ANOVA, Dunnett’s test versus mPPARα).

Histological analyses were carried out with H&E straining of the tibia from mice exposed for 14 weeks (**Figure 8A**). Effects of PFOA on trabecular histology were not apparent. We examined osteoblast activity by measuring serum Procollagen type I N-terminal propeptide (PINP) (**Figure 8B**). We took advantage of serum samples from a second PFOA exposure group (PFOA High, serum concentration of 99 µg/ml (Nielsen et al., 2025)). Importantly, in the overall ANOVA genotype remained significant (p=0.0038). In PPARα Null and mPPARα mice, the PFOA low exposure showed reduced serum PINP compared to Vh and PFOA high exposure resulted in significantly lower serum PINP compared to Vh. PFOA had no effect on serum PINP in hPPARα mice, however the serum concentration in all treatment groups for hPPARα mice was equivalent to the lowest concentrations in PPARα Null and mPPARα mice. mRNA expression of osteoblast biomarker genes showed a significant reduction in *Osx* expression in PPAR⍰ null and hPPAR⍰ mice compared to Vh (**Figure S5**).

**Figure 8.**
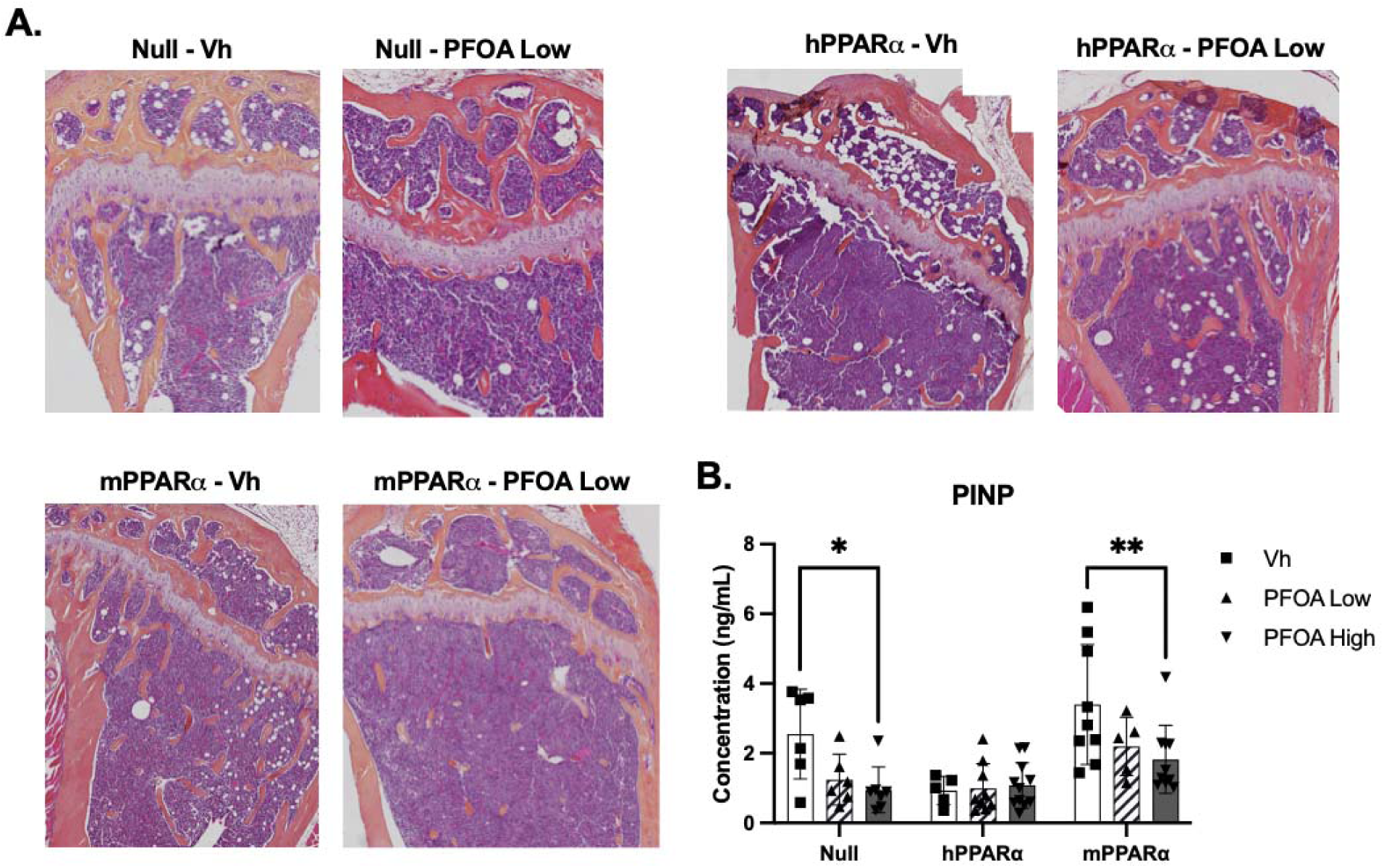
Analyses of bone formation for male and female mice after 14-week PFOA exposure. Samples were analyzed from the 14-week exposure described in Figure 1; mice were 20-weeks of age. **(A)** Representative (i.e. based on median PINP concentration for each genotype and treatment) images of 5 μm slices of distal femur stained with hematoxylin and eosin (4× magnification). **(B)** Quantification of serum PINP for males and females combined (*n* = 6–12). Samples with CV% > 30 were excluded (8 in total). Data points are from individual mice, with the mean indicated the bar and the SD indicated by the error bars. \**p <* 0.05, \*\**p <* 0.01 (ANOVA, Tukey’s multiple comparisons test within genotype).

Histological analyses were carried out for osteoclasts with Trap straining of the tibia from mice exposed for 14 weeks (**Figure 9A**). Trap staining was not obviously impacted by genotype or PFOA exposure. In the overall ANOVA for CTX serum concentration neither genotype nor treatment status were significant, p=0.126 and p=0.896, respectively. Similarly, there were no significant effects on serum C-terminal telopeptides of type I collagen (CTX-I) (**Figure 9B**). Furthermore, expression of biomarker genes of osteoclasts, *Acp5* and *Ctsk*, revealed no differences or significance of treatment or genotype (**Figure S6)**.

**Figure 9.**
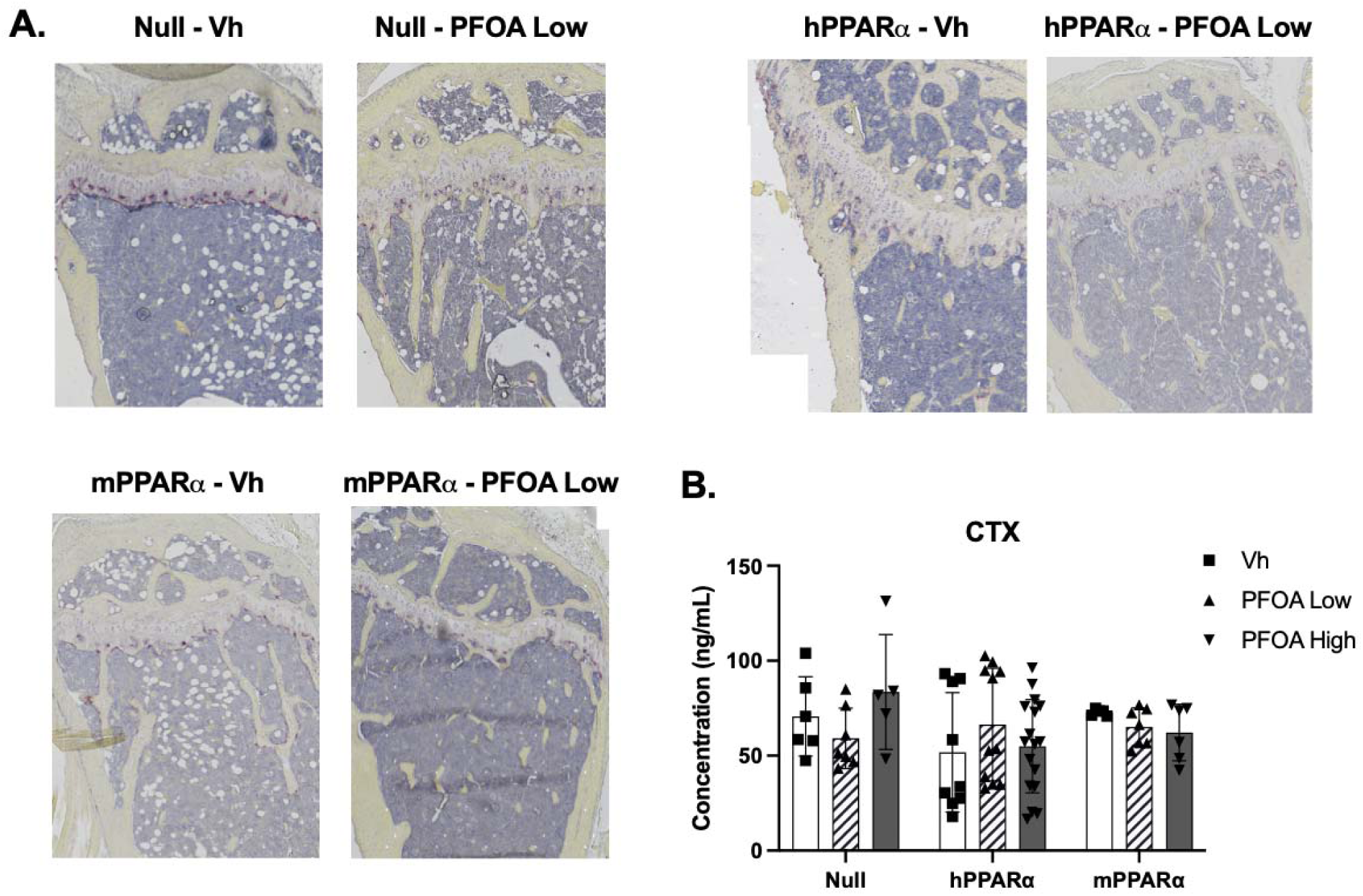
Analyses of bone resorption for male and female mice after 14-week PFOA exposure. Samples were analyzed from the 14-week exposure described in Figure 1; mice were 20-weeks of age. **(A)** Representative (i.e. based on median PINP concentration for each genotype and treatment) images of 5 μm slices of distal femur (*n* = 4–6) stained for TRAP (red shading) (10× magnification). Scale bar = 100 μm. **(B)** Quantification of serum CTX for males and females combined (*n* = 6–12). 1 outlier identified and removed in ROUT analysis. Samples with CV% > 30 were excluded (2 in total). Data points are from individual mice, with the mean indicated the bar and the SD indicated by the error bars. \**p <* 0.05, \*\**p <* 0.01 (ANOVA, Tukey’s multiple comparisons test within genotype).

## Discussion

Epidemiological evidence supports the hypothesis that PFAS negatively impacts bone quality. Studies on the effect of PFOA on bone health in animal models have produced varying results, with none carried out in mice expressing hPPARα. Therefore, the goal of this study was to investigate the effect of PFOA on bone quality in mice expressing human PPARα and being fed a human relevant diet. It was clear from analyses of the liver that a biologically effective dose of PFOA was achieved and that effects were modified by PFAS genotype. PFOA had little effect on bone structure. This contrasted with the strong effect of PPARα genotype on trabecular and cortical structure. However, previously reported effects of PFOA acting on bone were supported by analysis of markers of osteoblast activity, with PFOA exposure resulting in lower serum PINP and expression of *Osx* in bone. The results bolster the important role that PPARα plays in regulating bone quality and may support that PFOA can influence bone biology.

It is rare for analyses of PFOA-driven effects to include analyses at multiple time points or at a chronic time point; hence, we began with analyzing effects of PFOA in liver following 6-and 14-weeks exposures, which 1) generated similar PFOA serum concentrations and 2) occurred in the context of consumption of an American-type diet. PFOA exposure significantly increased liver weight and triglycerides across genotypes at 6 weeks. Notably, liver triglycerides were significantly increased in hPPARα and mPPARα mice but not in PPARα null mice. The overall hepatic triglyceride concentrations were higher at 14 weeks than 6 weeks. This is consistent with other studies of chronic exposure to a high (45%) fat diet (Patsouris et al., 2006). PFOA exposure did not further increase the concentrations of triglycerides after 14 weeks of exposure. The development of hepatosteatosis is consistent with previous evidence in mPPARα, hPPARα, and PPARα null mice (Nakamura et al., 2009; Schlezinger, Puckett, Oliver, et al., 2020; Su et al., 2022).

At both 6 and 14 weeks PFOA exposure, with serum PFOA concentrations of ≈ 50 µg/mL, exposure only significantly increased PPARα activation in mPPARα mice. Given the relatively low dose exposure to PFOA and the lower activity of hPPARα these results are not unexpected (Nakamura et al., 2009).

CAR and PXR were more strongly activated in hPPARα and PPARα null mice than in mPPARα mice. This is consistent with negative cross-talk between these nuclear receptors (Corton et al., 2014; Rosen et al., 2017; Schlezinger, Puckett, Oliver, et al., 2020). Importantly, after 14 weeks, PFOA activated CAR only in PPARα null mice, with only moderate activation in hPPARα - however this activation is substantially increased from 6 weeks from the range of 5-10 up to 20-40. Moreover, PXR activation at 14 weeks is markedly reduced with relative expression decreasing from nearly 300 to 10. These results indicate a greater activation of PXR during shorter exposures and of CAR during longer exposures. Our results strengthen the evidence for crosstalk between these nuclear receptors. However, it is not yet clear what the potential toxicological significance of the changing ratio of CAR to PXR activation over time is.

PPARα genotype was significant across all ANOVAs run for both cortical and trabecular bone parameters at both 6 and 14 weeks, emphasizing the importance of PPARα in regulating bone quality. Recently, it was reported that whole body knockout of PPARα led to 1) a higher total trabecular volume accompanied by a higher trabeculae number and 2) a larger total cortical area that was accompanied by a higher marrow area resulting in a lower bone:total area in males at 20 weeks of age compared to mPPARα mice (Chougule et al., 2023). In contrast, osteocyte-specific knockout of PPARα had no effect on bone architecture, suggesting that PPARα may have a lesser role to play in the most mature of bone cells. We observed that the greatest differences in trabecular bone structure across genotypes at 9 weeks of age, which were attenuated at 20 weeks of age. Trabecular thickness was greater and trabecular spacing was lesser in PPARα Null mice than mPPARα mice. Differences in cortical bone were more consistent between age groups, with PPARα Null mice have lower cortical bone volume to total volume accompanied by a lower cortical thickness, in particular. We also observed that PPARα Null mice had lower cortical tissue mineral density at both ages (an endpoint not measured in the Chougule et al. study). Importantly, both studies reported that PPARα genotype had significant influence on both trabecular and cortical bone architecture. The differences in the specific architectural changes may result from the fact that in our study mice were fed an American style diet (with 34-37% fat), while the other study used standard rodent chow, and fat content of the diet can have significant effects on bone architecture (Patsch et al., 2011).

However, there was no effect of PFOA exposure on either trabecular or cortical bone health parameters. Importantly, the 129/Sv background is not commonly used in studies of bone architecture and may limit detection of changes (Beamer et al., 1996; Bouxsein et al., 2005; Friedman et al., 2024; Ishimori et al., 2006). While 129/Sv mice have higher than average bone mineral density and cortical bone area their trabecular bone volume is low. As such, while we hypothesized that PFOA exposure would negatively impact bone quality the low dose exposure and the potential for limited detectability likely contributed to this null presenting finding. Furthermore, the micro-CT scanning parameters had to be altered for the trabecular structure for these mice, emphasizing the differences by strain.

Previous work investigating the role of PFAS on osteoblasts and osteoclasts emphasize its role in increasing resorptive activity with more variable results for their effect on formation (Di Nisio, De Rocco Ponce, et al., 2020; Garmo et al., 2025b; Koskela et al., 2016; Liu et al., 2019; Pan et al., 2019b; Wang et al., 2024b; S. Yang et al., 2026). Our results emphasize PFOA’s role in impacting osteoblast activity particularly for PPARα null and mPPARα mice, where PINP serum concentration was numerically reduced by PFOA in low exposure and significantly reduced by PFOA high exposure compared to the Vh. Interestingly, hPPARα mice had very low overall serum PINP concentrations being equal to or less than the lowest serum PINP concentration for PPARα null and mPPARα mice. Our data also show no impact of genotype or treatment status on CTX concentration. This contrasts with other studies. For instance, using bone marrow from mPPARα, hPPARα and PPARα null mice on the C57BL/6J background (which we backcrossed from the original hPPARα mice on the 129/sv background), it was shown that osteoclast differentiation was greatest in hPPARα bone marrow macrophages, followed by mPPARα macrophages and highly limited in PPARα Null macrophages (Garmo et al., 2025b). Further, a PFAS mixture further stimulated osteoclastogenesis in only the mPPARα mice. However, we have learned from our own studies that examining effects of toxicants on isolated bone forming or bone resorbing cells may not accurately represent what happens in vivo given the loss of the essential cross talk between these cell types (Watt et al., 2018). We also hypothesize that the genetic background may influence the bone outcomes.

Overall, our data supports the importance of PPARα genotype in determining trabecular and cortical bone parameters and osteoblast function. While PFOA significantly impacted the activation of nuclear receptors (CAR, PXR, and PPARα) in liver, it had little effect on bone architecture and quality. However, that PFOA reduced a biomarker of osteoblast activity is another piece of evidence that supports a potential cause-and-effect relationship in the epidemiological evidence showing associations between PFAS exposure and loss of bone quality.

## Acknowledgements

This work was supported by funding from the National Institutes of Health. The original experiment studies were supported by R01 ES027813 (Thomas Webster, PI). The bone analyses were supported by R03 ES034093 (Jennifer Schlezinger, PI). We would like to thank Lauren Payne and Artur Kuchumov for their assistance the gene expression analyses. Sydney Holder of the Boston University Micro-Computed Tomography Imaging Core Facility provided essential assistance in the microCT analyses. Lang Ma of the MD Anderson Cancer Center Bone Histomorphometry Core Laboratory provided essential assistance in the bone histology analyses.

## Notes

### Competing Interest Statement

The authors have declared no competing interest.

### Summary of Updates

To update the introduction to include a new summary table in the supplemental that includes all the existing in vivo studies carried out on PFAS and bone health

