## Supplemental Data for "PPARα genotype impacts bone quality and perfluorooctanoic acid-induced effects"

Carpenter, M.O., Schlezinger, J.J.

Table of Contents

| Table S1. | *In vivo* studies of PFAS-induced effects on bone | 2 |
| --- | --- | --- |
| Table S2. | Components of the “What we eat in America” diets (Research Diets D18110214) | 3 |
| Table S3. | Mouse and human primer sequences for reverse transcriptase qPCR | 4 |
| Table S4. | ANOVA results for Trabecular bone structure after 6 and 14 week exposure | 5 |
| Table S5. | ANOVA results for Cortical bone structure after 6 and 14 week exposure | 5 |
| Figure S1. | Micro-CT analysis of Male trabecular bone structure after 6 week exposure | 6 |
| Figure S2. | Micro-CT analysis of Male trabecular bone structure after 14 week exposure | 7 |
| Figure S3. | Micro-CT analysis of Male cortical bone structure after 6 week exposure | 8 |
| Figure S4. | Micro-CT analysis of Male cortical bone structure after 14 week exposure | 9 |
| Figure S5. | mRNA expression of osteoblast biomarker genes after 14 week exposure | 10 |
| Figure S6. | mRNA expression of osteoclast biomarker genes after 14 week exposure | 11 |

**Table S1. *In vivo* studies of PFAS-induced effects on bone**

| **Study** | **Background/ Sex** | **Timing of Exposure** | **Age at Analysis** | **PFAS** | **Dosing** | **Serum Conc.(μg/mL)** | **Endpoint** |
| --- | --- | --- | --- | --- | --- | --- | --- |
| **Lau et al., 2006** | CD-1  Dam Exposed | GD1-GD17 | GD18 | PFOA | Gavage, 1 mg/kg/day | ≈ 25 (maternal) | Reduced calavarial ossification |
| **Koskela et al., 2016** | C57BL/6/Bkl Dam Exposed | GD1-GD21 | 17 months | PFOA | Food, 0.3 mg/kg/day | Not  measured* | Increased femoral and tibial periosteal and medullary area |
|  |  |  |  |  |  |  | Reduced femoral and tibial tissue mineral density |
| **Feng et al., 2024** | C57BL6/J Dam Exposed | GD1-PND21 | 3 weeks | 6:2 Cl-PFESA | Drinking water, 5.7 mg/L | Not measured | In males, shortened femur length, reduced trabecular and cortical bone quality parameters |
| **Wang et al., 2025** | BALB/c Male | 3-7 weeks of age | 7 weeks | PFOA | Gavage,  2 mg/kg/day | Not measured | Shortened femur length, reduced trabecular bone quality  parameters |
|  |  |  |  | HFPO-DA | Gavage,  4 mg/kg/day |  |  |
|  |  |  |  | FO3TDA | Gavage, 0.4 mg/kg/day |  |  |
| **Garmo et al., 2025** | FVB/N  Male | 12 weeks, ages not reported | | HFPO-DA | Drinking  water, 2 mg/L | 0.14 | Reduced cortical and trabecular bone volume in tibia |
|  |  |  |  | PFHxS | Drinking  water, 2 mg/L | 24 |  |
| **Current Study** | 129/Sv  Male and Female | 6-20 weeks of age | 20 weeks | PFOA | Drinking  water, 1.4 mg/L | 50 | No effects on femoral bone structure |
|  |  |  |  |  | Drinking  water, 6.2 mg/L | 100 | Reduced serum PINP in PPARα Null and mPPARα mice |

* PFOA concentration in bone was elevated over control by 5.8-fold.

Abbreviations: HFPO-DA - perfluoro (2-methyl-3-oxhexanoic) acid; PFOA - perfluorooctanoic acid; PFO3TDA - perfluoro-3,6,9-trioxyundecanoic acid; PFHxS - perfluorohexane sulfonic acid; 6:2 Cl-PFESA - 2-((6-chloro-1,1,2,2,3,3,4,4,5,5,6,6-dodecafluorohexyl)oxy)-1,1,2,2-tetrafluoroethanesulfonic acid)

### Table S1. Components of the “What we eat in America” diets.

|  | Adolescent  (Research Diets D18110214) | | Adult  (Research Diets D20031203) | |
| --- | --- | --- | --- | --- |
| Macronutrient | % Grams | % kcal | % Grams | % kcal |
| Protein | 16 | 14.7 | 18 | 16 |
| Carbohydrate | 57 | 51.8 | 54 | 47 |
| Fat | 16 | 33.5 | 19 | 37 |
| Fiber | 5.4 |  | 5.5 |  |
| Cholesterol | 0.05 |  | 0.13 |  |
| Kcal/gm | 4.4 |  | 4.5 |  |
| Ingredient | grams | kcal | Grams | kcal |
| Casein | 146 | 584 | 158.5 | 634 |
| L-cysteine | 3 | 12 | 4.5 | 18 |
| Corn Starch | 177 | 707 | 163 | 652 |
| Maltodextrin 10 | 100 | 400 | 100 | 400 |
| Sucrose | 238.67 | 955 | 201.67 | 807 |
| Cellulose | 40 | 0 | 40 | 0 |
| Inulin | 10 | 15 | 10 | 15 |
| Soybean Oil | 25 | 225 | 25 | 225 |
| Lard | 75.6 | 680 | 92 | 828 |
| Butter | 50.4 | 454 | 50.4 | 454 |
| Mineral Mix S10026 | 10 |  | 10 | 0 |
| Dicalcium Phosphate | 13 |  | 13 | 0 |
| Calcium Carbonate | 5.5 |  | 5.5 | 0 |
| Potassium Citrate Monohydrate | 16.5 |  | 16.5 | 0 |
| Vitamin Mix V10001 | 10 | 40 | 10 | 40 |
| Choline Bitartrate | 2 |  | 2 | 0 |
| Cholesterol | 0.2943 |  | 1 | 0 |
| FD&C Yellow Dye #5 | 0.025 |  | 0 | 0 |
| FD&C Red Dye #40 | 0.025 |  | 0 | 0 |
| FD&C Blue Dye #1 | 0 |  | 0.05 |  |
| Total | 922.82 | 4072 | 903.12 | 4072 |

Table S2. Primer sequences for reverse transcriptase qPCR.

| Gene  Symbol | FORWARD | REVERSE | Annealing Temp. °C |
| --- | --- | --- | --- |
| R18s | GTAACCCGTTGAACCCCATT | CCATCCAATCGGTAGTAGCG | 55 |
| Acox | AGCGAGCCAGAGCCCCAG | TCAGGCAGCTCACTCAGG | 59 |
| Cyp2b10 | AAAGTCCCGTGGCAACTTCC | TCCCAGGTGCACTGTGAACA | 61 |
| Cyp3a11 | ACAAACAAGCAGGGATGGAC | GGTAGAGGAGCACCAAGCTG | 57 |
| Osx/Sp7 | GTCAAGAGTCTTAGCCAAACTC | AAATGATGTGAGGCCAGATGG | 55 |
| Dmp1 | CAACTGGCTTTTCTGCGGCAA | TGGGTGCGCTGATGTTTGCT | 55 |
| Trap5/Acp5 | CCATTGTTAGCCACATACGG | ACTCAGCACATAGCCCACAC | 55 |
| Ctsk | GTTGTATGTATAACGCCACGGC | CTTTCTCGTTCCCCACAGGA | 60 |

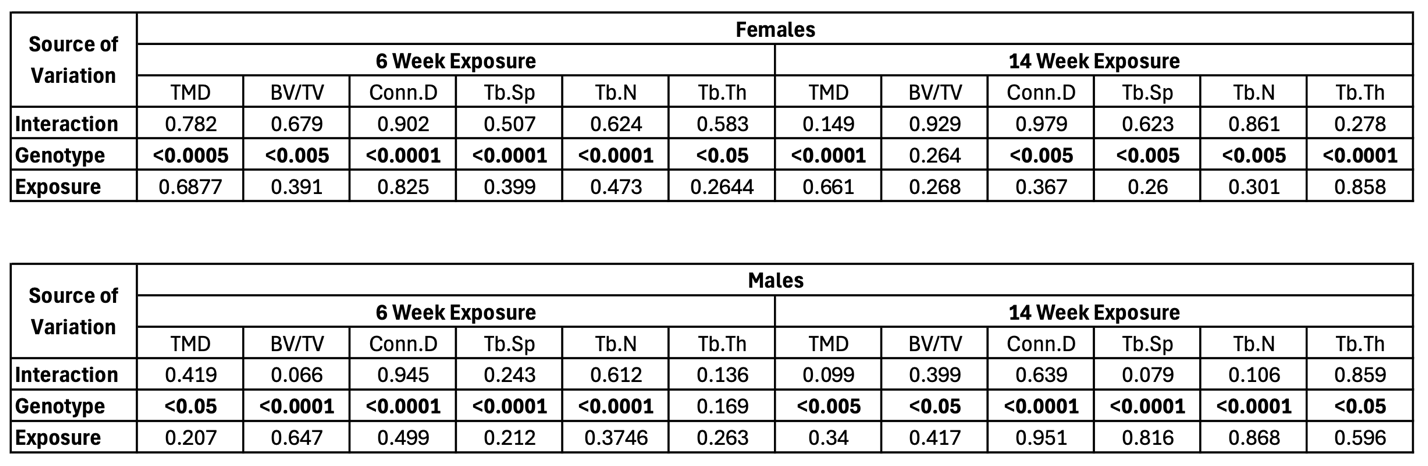

Table S3. ANOVA results for Trabecular Bone structure for Males and Females 9 and 20 weeks of Age.

Two-way ANOVA carried out for each trabecular bone parameter. N=6-9. Significant p-values bolded.

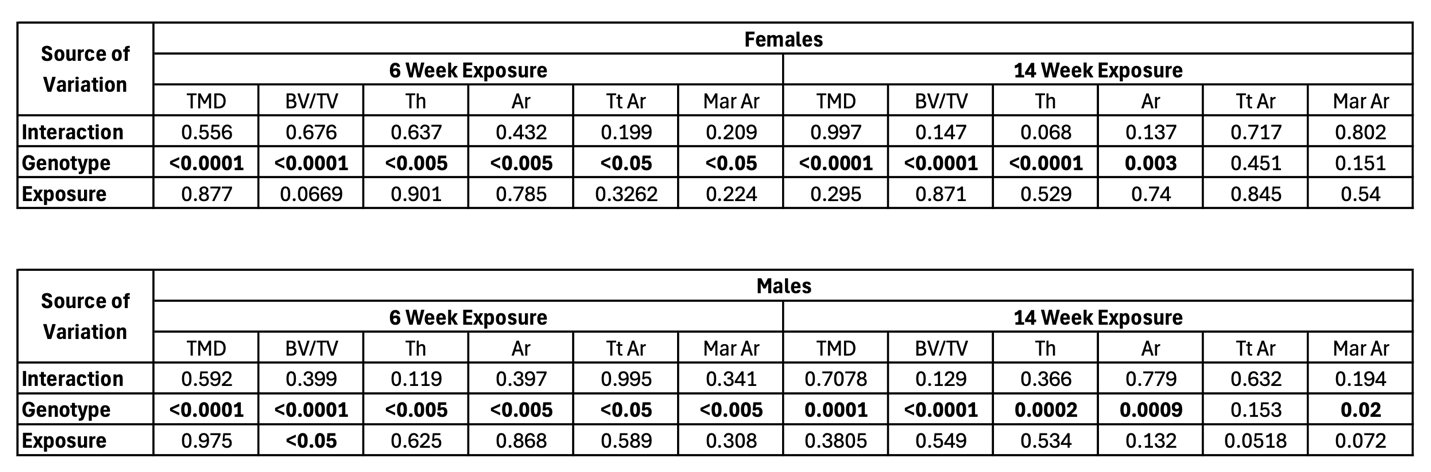

### Table S4. ANOVA results for Cortical bone structure data for Males and Females 9 and 20 weeks of Age.

Two-way ANOVA carried out for each trabecular bone parameter. N=6-9. Significant p-values bolded.

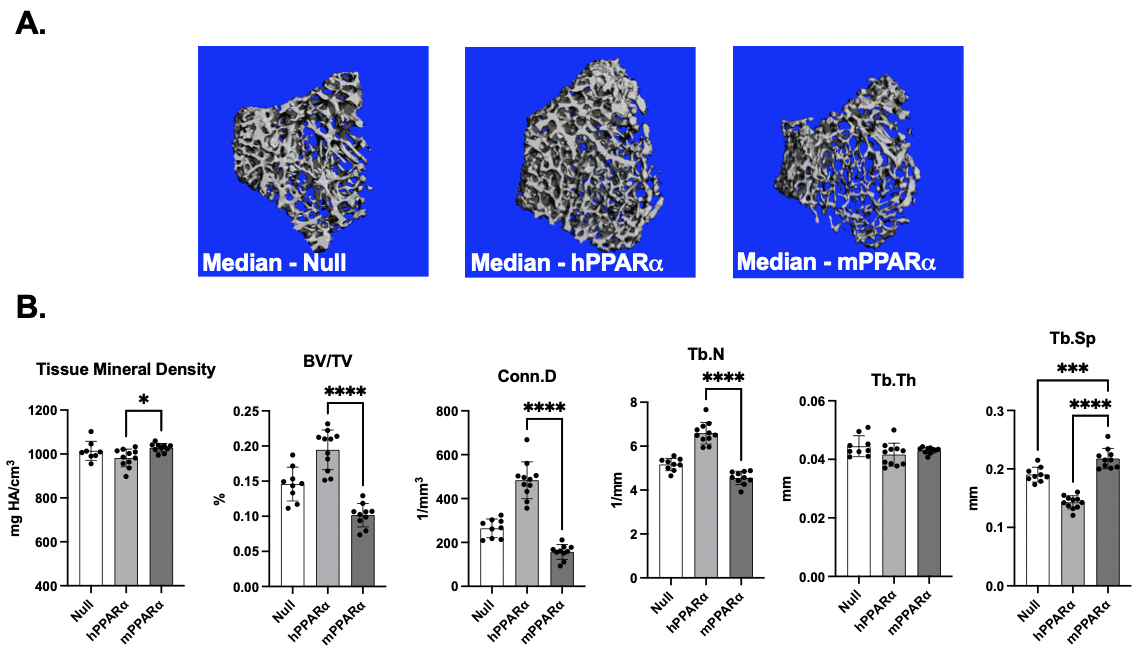

### Figure S1. Micro-CT analysis of femur trabecular bone structure for male mice after 6-week PFOA exposure.

Samples analyzed from the 6-week exposure described in Figure 1, mice were 9-weeks of age. **(A)** Representative (i.e., bones with the median connective density for each genotype) micro-CT images of distal metaphysis of male mice. **(B)** Trabecular bone parameters of male mice. Data points are from individual mice, with the mean indicated the bar and the SD indicated by the error bars. N = 6-9. *p<0.05, ** p<0.01, *** p<0.001, ****p<0.0001 (ANOVA, Dunnett’s test versus mPPARα).

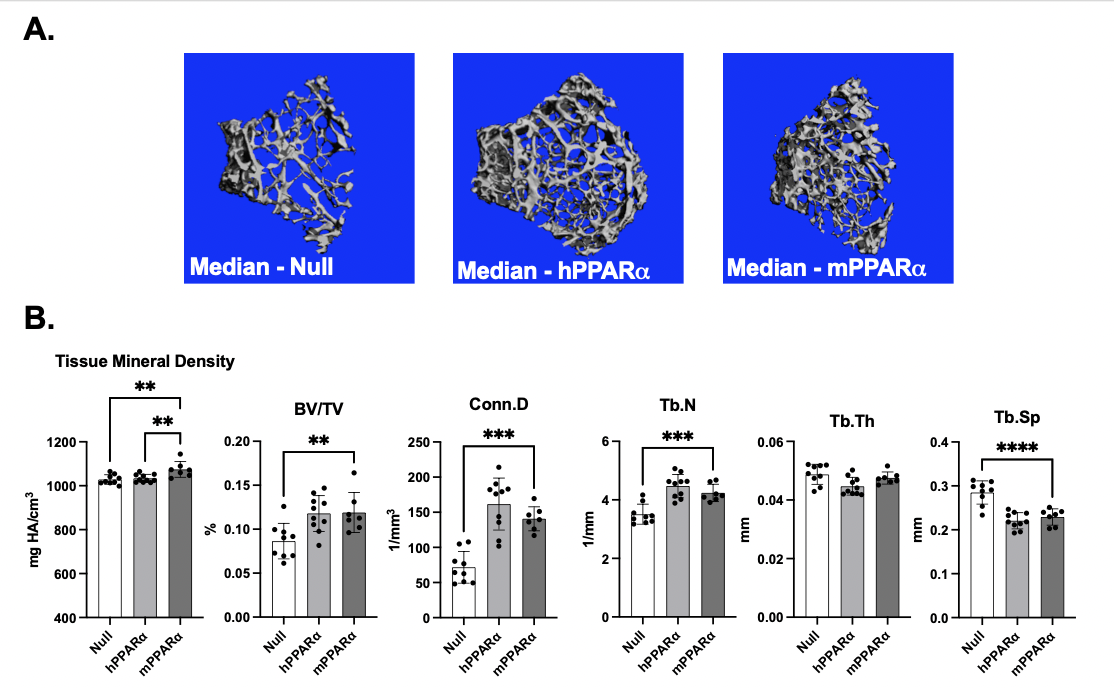

Figure S2. Micro-CT analysis of femur trabecular bone structure for male mice after a 14-week PFOA exposure.

Samples analyzed from the 14-week exposure described in Figure 1, mice were 20 weeks of age. **(A)** Representative (i.e., bones with the median connective density for each genotype) micro-CT images of distal metaphysis of male mice. **(B)** Trabecular bone parameters of male mice. Data points are from individual mice, with the mean indicated the bar and the SD indicated by the error bars. N = 6-9. *p<0.05, ** p<0.01, *** p<0.001, ****p<0.0001 (ANOVA, Dunnett’s test versus mPPARα).

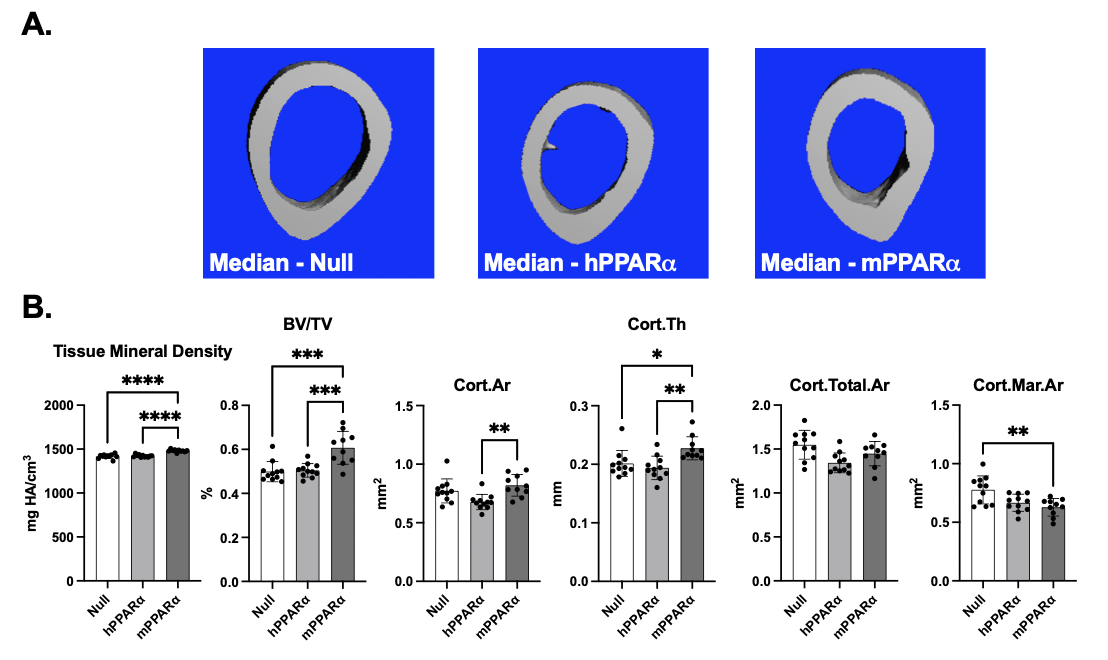

### Figure S3. Micro-CT analysis of femur cortical bone structure for male mice after 6-week PFOA exposure.

Samples analyzed from 6-week exposure described in Figure 1, mice were 9-weeks of age. **(A)** Representative (i.e. bones with median cortical thickness for each genotype) micro-CT images of mid-diaphysis of male mice. **(B)** Cortical bone parameters of male mice. BV/TV: bone volume to total volume ratio, Cort.Ar: total cortical area, Cor.Th: cortical thickness, Cort.Total.Area: total bone area, Cort.Mar.Ar: Cortical marrow area. Data points are from individual mice, with the mean indicated the bar and the SD indicated by the error bars. N = 6-9. *p<0.05, ** p<0.01, *** p<0.001, ****p<0.0001 (ANOVA, Dunnett’s test versus mPPARα).

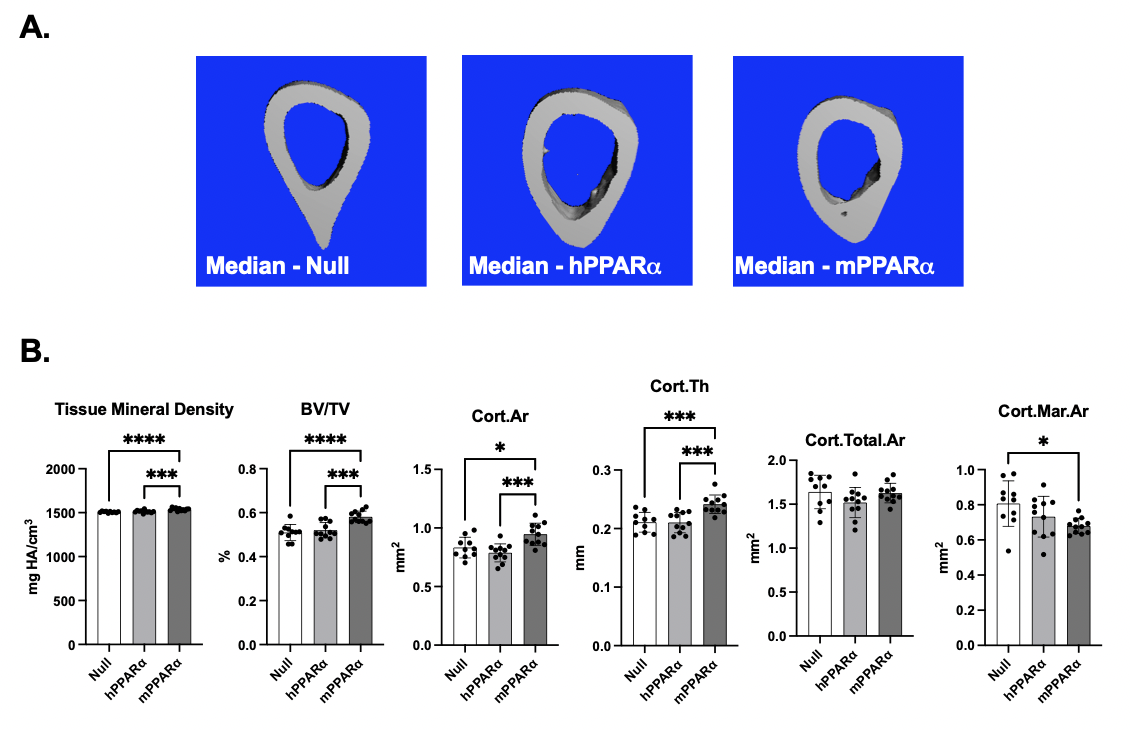

### Figure S4. Micro-CT analysis of femur cortical bone structure for male mice after 14-week PFOA exposure.

Samples analyzed from the 14-week exposure described in Figure 1, mice were 20-weeks of age. **(A)** Representative (i.e. bones with median cortical thickness for each genotype) micro-CT images of mid-diaphysis of male mice. **(B)** Cortical bone parameters of male mice. BV/TV: bone volume to total volume ratio, Cort.Ar: total cortical area, Cor.Th: cortical thickness, Cort.Total.Area: total bone area, Cort.Mar.Ar: Cortical marrow area. Data points are from individual mice, with the mean indicated by the bar and the SD indicated by the error bars. N = 6-9. *p<0.05, ** p<0.01, *** p<0.001, ****p<0.0001 (ANOVA, Dunnett’s test versus mPPARα).

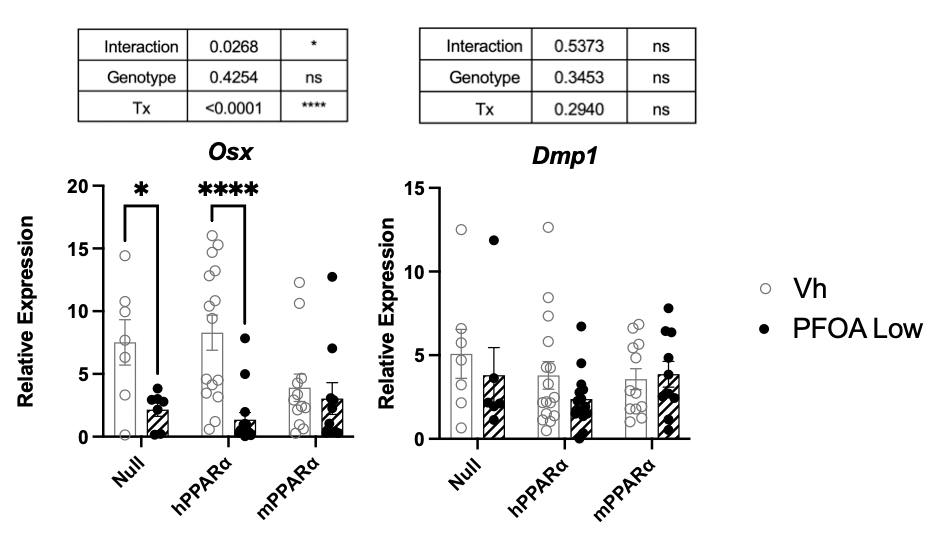

### Figure S5. mRNA expression of osteoblast biomarker genes (*Osx and Dmp1*) in male and female mice after 14-week PFOA exposure.

Samples analyzed from 14-week exposure described in Figure 1, mice were 20 weeks of age. mRNA expression of selected biomarkers of osteoblast function from whole tibia. Data points are from individual mice with the mean indicated by the bar and the SD indicated by the error bars. N= 7-16. *p<0.05, ** p<0.01, *** p<0.001 (ANOVA, Sidaks test versus Vh).

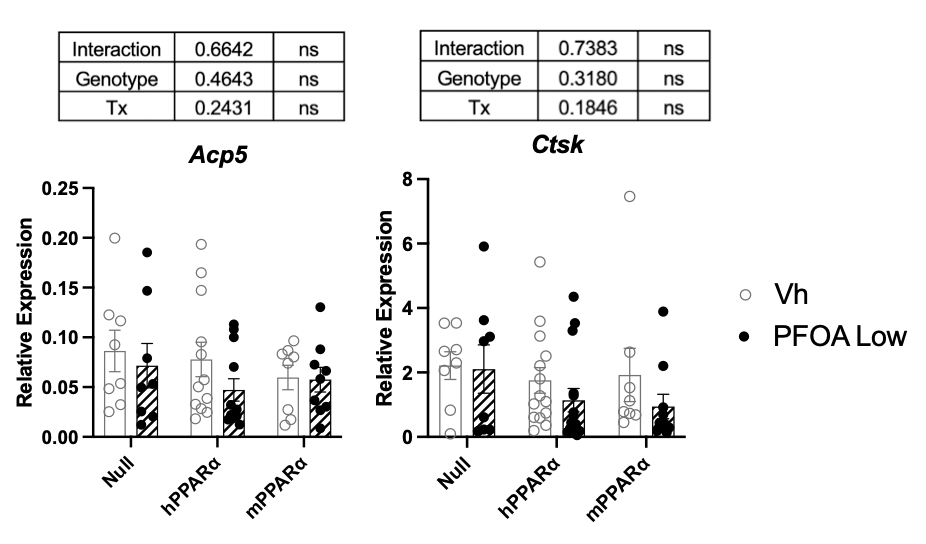

### Figure S6. mRNA expression of osteoclast biomarker genes (*Acp5 and Ctsk*) in male and female mice after 14-week PFOA exposure.

Samples analyzed from 14-week exposure described in Figure 1, mice were 20 weeks of age. mRNA expression of selected biomarkers of osteoclast function from whole tibia. Data points are from individual mice with the mean indicated by the bar and the SD indicated by the error bars. N= 7-16. *p<0.05, ** p<0.01, *** p<0.001 (ANOVA, Sidaks test versus Vh).
